# A Hydrophobic Microenvironment Significantly Influences the Reactivity of the Catalytically Relevant Thiols of the Na^+^/K^+^-ATPase

**DOI:** 10.1101/2022.10.30.514419

**Authors:** Titilayo Ibironke Ologunagba, Tolulope Ayoola Ojo, Ige Joseph Kade

## Abstract

The transmembrane protein responsible for the electrogenic transport of Na^+^ and K^+^ across the plasma membrane, the Na^+^/K^+^-ATPase, highly vulnerable redox modulations and thiol modifying agents due to the presence of thiol groups at the nucleotide and cationic sites. However, reports have demonstrated a preferential interaction of these protein thiols with oxidizing agents. The reactivity of protein thiols is strongly linked with the nature of the microenvironment of these thiols, hence, the present study sought to experimentally elucidate key features of the microenvironment of the catalytically relevant thiols at the substrate-binding sites of this crucial enzyme. Two thiol modifiers with similar thiol-reactive mechanism, but different molecular properties, iodoacetamide (IA) and N-acetyl-4-phenyliodoacetamide (APIAM), were employed. It was observed that while both compounds demonstrated excellent thiol-oxidizing properties in the chemical model, only APIAM had an inhibitory effect on the activity of the Na^+^/K^+^-ATPase. The involvement of the catalytically relevant thiols at the nucleotide and cation-binding sites of the enzyme in APIAM-mediated inhibition was confirmed by the protective effect of preincubating the reaction system with dithiothreitol (DTT). The findings from this study suggest that the catalytically relevant thiols of this enzyme are likely buried in a hydrophobic microenvironment. This could be a part of the protective measure of nature for these vulnerable protein thiols. Further details from our findings can be explored in the therapeutic management of diseases for which a dysfunction in the Na^+^/K^+^-ATPase have been identified.

**Highlights:**

- The transmembrane Na^+^/K^+^-ATPase has well-defined substrate-binding domains exposed to both aqueous microenvironment and buried within the hydrophobic transmembrane microenvironment
- These microenvironments influence vulnerability of the critical thiols of the enzyme to oxidative assault
- These thiols are likely buried in the hydrophobic core of the enzyme, thus selecting its susceptibility to thiol modfying agents

## INTRODUCTION

The maintenance of Na^+^ and K^+^ gradients across the plasma membrane, primarily coordinated by the Na^+^/K^+^-ATPase, is critical to cell survival. This electrochemical gradient is necessary for driving several physiological processes such as, Na^+^-coupled transport of nutrients and amino acids into cells, osmotic balance, cell volume regulation, and for maintenance and restoration of the resting membrane potential in excitable cells (Geering, 1997; Pavlov and Sokolov, 2000). First described by Danish researcher Jens Skou in 1957 (Skou, 1957), the Na^+^/K^+^-ATPase (NKA) is said to account for 23% of the ATP utilization in a resting human (Kaplan, 2002). Beyond its electrogenic function, this enzyme also serves as a scaffolding protein for intracellular signaling, being complexed with the non-receptor Src kinase (tyrosine kinase), caveolin and epidermal growth factor receptor (EGFR) to form a signalosome in the caveolar of cytoplasmic membrane (Yosef et al. 2016; Liu et al. 2011). The regulation of intracellular signaling cascades as well as enzyme hydrolytic and transport function of the NKA by oxidative oscillations is widely reported (Kade et al., 2008; Omotayo et al., 2011; 2015; Rauchova et al., 1999). More so, reports have consistently associated a modulation in the activity of this enzyme to different pathological conditions (Bejcek et al., 2021; Shrivastava et al., 2015; Dichiara et al., 2017). In fact, there is a growing interest in the exploration of the structure and physiological function of this crucial enzyme in the therapeutic management of a number of oxidative stress-linked pathologies (Aydemir-Koksoy et al., 2021; Chen et al., 2019; Khajah et al., 2018; Prassas et al., 2011). Hence, an understanding of its structure-function relationship thus gives further insight into its vulnerability to altered redox status.

The functional Na^+^/K^+^-ATPase complex is composed of three subunits, an α-subunit, a β-subunit and a FXYD (γ)-subunit. The α-subunit is a large polypeptide containing 10-transmembrane helices which form the binding sites for Na^+^, K^+^ and the specific inhibitor, ouabain (Kaplan, 2002; Blanco et al., 1998; Malik et al., 1996). It also has three cytoplasmic domains,: the nucleotide-binding domain (functioning as a kinase), the phosphorylation domain (functioning as the substrate), and the actuator domain (functioning as a phosphatase) (Shrivastava et al., 2018; Clausen et al., 2017; Toyoshima et al., 2011; Tidow et al., 2010). The β-subunit is responsible for the targeting and insertion of the α-subunit in the membrane and also modulates cation binding affinity and K^+^ occlusion (Toyoshima et al., 2011; Tidow et al., 2010; Barwe et al. 2007; Kaplan 2002), as well as cell adhesion and motility (Geering 2008). The FXYD unit is a small hydrophobic protein (Yoneda et al., 2020) that may not have a role in catalysis but in enzyme regulation, modulating Na^+^, K^+^, and ATP apparent affinity (Geering 2008; Beguin et al. 1997).

Individual subunits, domains or regions of this enzyme have been reported as targets of oxidative assaults due to the presence of conserved cysteine residues that are subject to glutathionylation in oxidative conditions (Yoneda et al., 2020). In fact, Petrushanko and coworkers earlier described the presence of four cysteine residues on the α-subunit which are subject to regulatory glutathionylation (Petrushanko et al., 2012). Furthermore, the group reported the presence of basal glutathionylation on the α-subunit which was not removed by dithiothreitol (DTT). It was suggested that the basally glutathionylated cysteine residues were located in a region of protein structure that was inaccessible to solvent (Mitkevich et al., 2016). The beta subunit also contains seven cysteine residues, six of which form three extracellular disulphide bonds, while one is predicted to be located within the membrane-panning region of the polypeptide (McDonough et al., 1990). The FXYD subunit also has two conserved cysteine residues in its C-terminus which are susceptible to thiol modification (Bibert et al., 2011). Taken together, it is apparent that the cysteine residues of the enzyme have clearly defined microenvironments that may be differentially accessible to different compounds.

The reactivity of these cysteine residues has been significantly related to the presence of other charged amino acid residues in their vicinity, which define their microenvironments. Furthermore, the Na^+^/K^+^-ATPase is an integral membrane protein which experiences two distinct environments defined by the membrane namely; a hydrophilic (extracellular/cytosolic) environment and a hydrophobic environment (within the membrane lipid bilayer) (Yoneda et al., 2020). This feature likely contributes to the availability of different domains of this protein for interaction with exogenous compounds. In fact, an earlier report has highlighted a differential sensitivity of the enzyme to two organoselenium-based compounds with significantly different molecular sizes, dicholesteroyl diselenide and diphenyl diselenide (Kade et al., 2009). While dicholesteroyl diselenide (DCDSe) inhibited the activity of another sulfhydryl protein, lactate dehydrogenase, to almost the same degree as diphenyl diselenide (DPDSe), it showed far less inhibitory effect against Na^+^/K^+^-ATPase. The authors suggested that the volume of the non-selenium moiety of DCDSe may pose an hinderance in its access to the thiols at the catalytic site of the Na^+^/K^+^-ATPase.

Apparently, there is likely a principle of selection operational at the substrate binding sites vis-a-viz the spatial location of the thiols of this crucial enzyme guiding their vulnerability to a variety of oxidative agents. The present study employs two thiol-reactive compounds, iodoacetamide and N-acetyl-4-phenyliodoacetamide, having different polarities to investigate the nature of the microenvironment of the catalytically relevant thiols of the Na^+^/K^+^-ATPase.

## MATERIALS AND METHODS

### Chemicals

Adenosinetriphosphate (ATP), iodoacetamide (IA), N-acetyl-4-phenyliodoacetamide (APIAM), dithiothreitol, ouabain, 5’5’-dithio-bis (2-nitrobenzoic) acid (DTNB) were obtained from Sigma (St. Louis, MO). All other chemicals which are of analytical grade were obtained from standard commercial suppliers.

### Animals

Male adult Wistar rats (200–250g) from our own breeding colony were used. Animals were kept in separate animal cages, on a 12-h light: 12-h dark cycle, at a room temperature of 22–24°C, and with free access to food and water. The animals were used according to standard guidelines on the Care and Use of Experimental Animal Resources.

### Preparation of synaptosomal fraction

Rats were decapitated under mild ether anaesthesia and the cerebral tissue (whole brain) was rapidly removed, placed on ice and weighed. The brain was immediately homogenized in cold 100mM Tris–HCl containing 300mM sucrose, pH 7.4 (1/10, w/v) with 10 up-and-down strokes at approximately 1200 rev/min in a Teflon–glass homogenizer. The homogenate was centrifuged for 10 min at 4000g to yield a pellet that was discarded and a low-speed supernatant(S1). The supernatant S1 was then centrifuged at 12000rpm for 10 minutes to yield a membrane and mitochondria-rich synaptosomal fraction as pellet and cytosolic fraction as supernatant. The supernatant was discarded and the pellet was reconstituted in 50mM Tris-HCl, pH 7.4 (1/5, w/v).

### Incubation system for sodium pump assay

Aliquot of the synaptosomal fraction was used for the assay of Na^+^/K^+^-ATPase activity. The reaction mixture contained 3mM MgCl_2_, 125mM NaCl, 20mM KCl and 50mM Tris–HCl, pH 7.4, iodoacetamide (final concentrations range of 0.1–1mM), N-acetyl-4-phenyliodoacetamide (final concentrations range of 0.1–1mM), with and without dithiothreitol (final concentration, 2 mM), and 100–180 mg protein in a final volume of 500 ml. The reaction was initiated by addition of ATP to a final concentration of 3.0mM. Controls were carried out under the same conditions with the addition of 0.1mM ouabain. The reaction mixture was incubated at 37°C for 30 min. At the end of the incubation period, tubes were assayed for sodium pump activity.

### Assay of sodium pump

The reaction system for the assay of the activity of cerebral Na^+^/K^+^-ATPase was essentially the same as described above under the section “incubation system for sodium pump assay”. However, at the end of the incubation time period (30–60 min), the reaction was stopped by addition of 5% trichloroacetic acid. Released inorganic phosphorous (Pi) was measured by the method of Fiske and Subbarow (1925). Na^+^/K^+^-ATPase activity was calculated by the difference between two assays (with and without ouabain). All the experiments were conducted at least three times and similar results were obtained. Protein was measured by the method of Lowry *et al*. (1951), using bovine serum albumin as standard. For all enzyme assays, incubation times and protein concentration were chosen to ensure the linearity of the reactions. All samples were run in duplicate. Controls with the addition of the enzyme preparation after mixing with trichloroaceticacid (TCA) were used to correct for non-enzymatic hydrolysis of substrates. Enzyme activity was expressed as nmol of phosphate (Pi) released min^-1^ mg protein^-1^.

### Oxidation of thiols

The rate of cysteine (CYS), glutathione (GSH) and dithiothreitol (DTT) oxidation was determined in the presence of 50mM Tris–HCl, varying pH and at pH 7.4 in the presence of IA [2mM] and APIAM [2mM]. The rate of thiol oxidation was evaluated by measuring the disappearance of –SH groups. Free –SH groups were determined according to Ellman (1959). Incubation at 37°C was initiated by the addition of DTT (final concentration, 1mM). Aliquots of the reaction mixture (100 ml) were checked (at 1h interval for 3h) for the amount of –SH groups at 412 nm by the addition of the colour reagent 5’5’-dithio-bis (2-nitrobenzoic) acid (DTNB).

### Statistical analysis

Results were analysed by appropriate analysis of variance (ANOVA) and this is indicated in text of results. Duncan’s Multiple Range Test was applied. Differences between groups were considered to be significant when p <0.05.

## RESULTS

### Chemical model: effect of thiol modifiers, iodoacetamide and N-acetyl-4-phenyliodoacetamide, on oxidation of thiols

The effect of thiol modifying agents, iodoacetamide (IA) [2mM] and N-acetyl-4-phenyliodoacetamide (APIAM) [1mM] on oxidation of a dithiol, dithiothreitol [1mM] is presented in Figure 1 (A and B). Herein, both thiol modifiers significantly oxidized the dithiol in a time-dependent fashion. This chemical model is a mild simulation of the potent oxidizing effect IA (panel A) and APIAM (panel B) following interaction thiols, and could give a prelude to their interaction in simple biological models.

**Figure 1:**
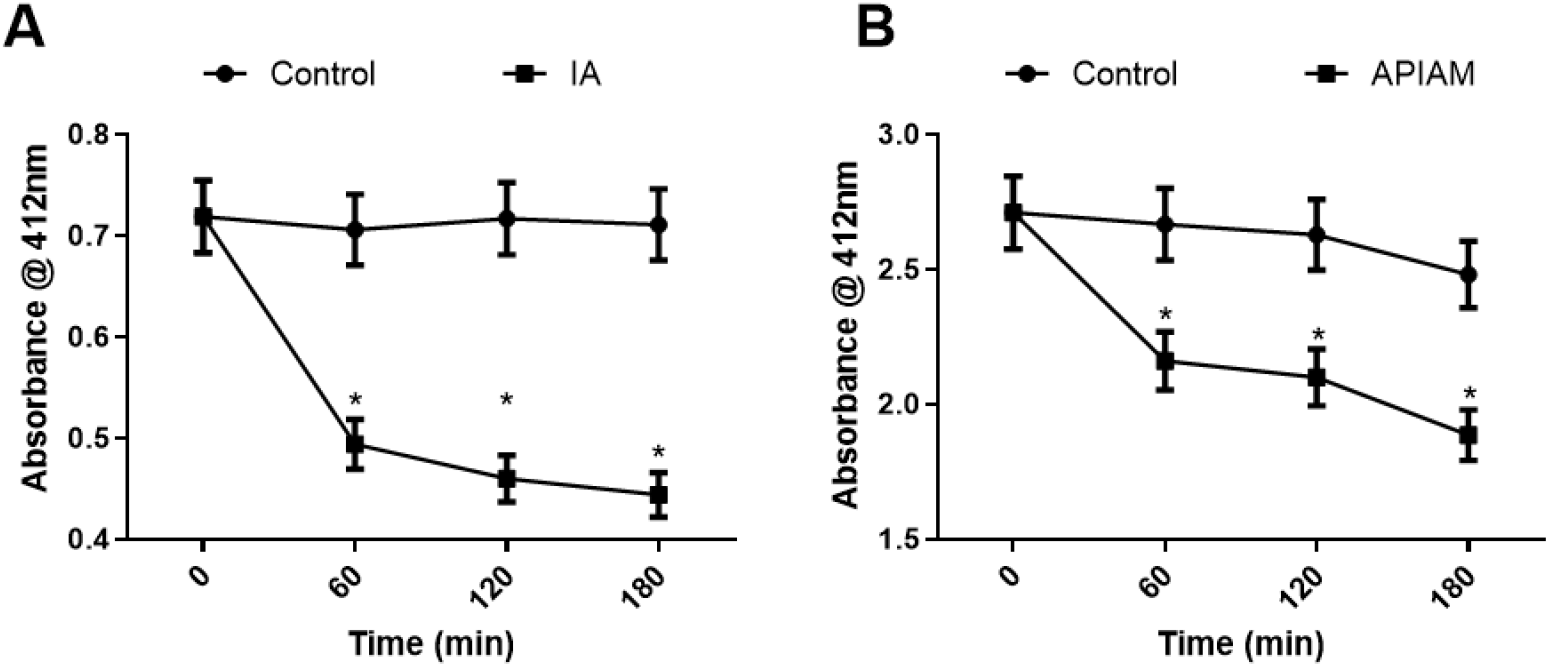
Chemical model of the effect of IA [2mM] (panel A) and APIAM [1mM] (panel B) on the rate of oxidation of DTT [1mM]. The rate of oxidation was evaluated at the indicated time-points. Data were the means of three independent experiments carried out on different days. Data were presented as mean ± SEM of at least three independent experiments carried out on different days. * indicates significantly lower than control (p < 0.05).

### Effect of thiol modifiers on the activity of the pump

The physiological relevance of the chemical model demonstrated above is shown by a simple biological model *in vitro* using membrane-rich synaptosomal fraction of the cerebral tissue of Wistar albino rats, to evaluate the effect of IA and APIAM on the activity of Na^+^/K^+^-ATPase at the ATP-binding site. One-way ANOVA analysis of the results showed that APIAM rather than IA, mediated inhibition of the sodium pump in a concentration-dependent manner, at the nucleotide-binding site (Figure 2, panel B). However, IA had no inhibitory effect on the activity of the electrogenic pump (panel A).

**Figure 2:**
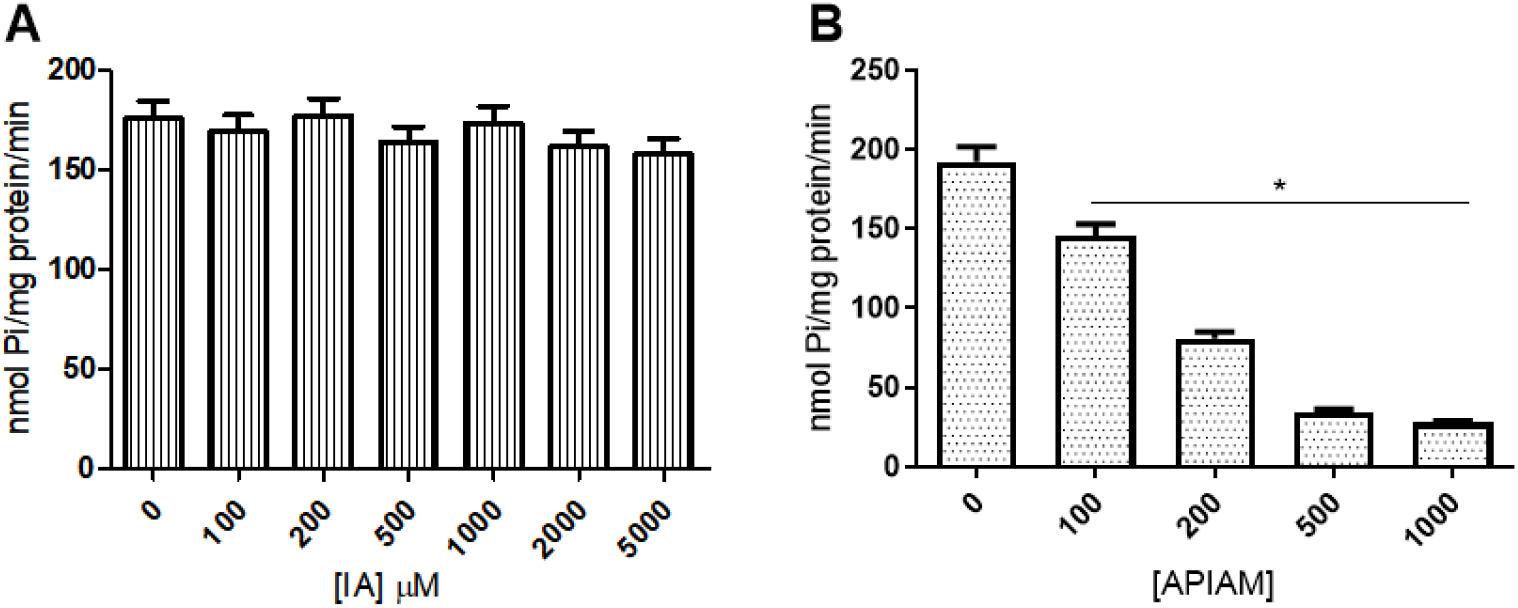
Effect of varying concentrations of IA (panel A) and APIAM (panel B) on the activity of the Na^+^/K^+^-ATPase at the nucleotide-binding site. Data were presented as mean ± SEM of at least three independent experiments carried out on different days. # indicates significantly lower than control (p < 0.05).

### Effect of exogenous thiol, dithiothreitol, on APIAM-mediated enzyme inactivation

Having demonstrated the thiol-oxidizing property of APIAM from the chemical model, it is rational to opine that its inhibition of the sulfhydryl Na^+^/K^+^-ATPase involves interaction with thiol groups at the nucleotide-binding site of this enzyme. Hence, the effect of pre- and post-incubation with exogenous thiol, dithiothreitol, on the inhibition mediated by APIAM was investigated. As presented in Figure 3, two-way ANOVA analysis of the results revealed that preincubation with DTT (panel A), significantly prevented the inhibition mediated by APIAM on the activity of the pump (p < 0.05). In contrast, post-incubation with DTT did not reverse APIAM-mediated enzyme inactivation (panel B).

**Figure 3:**
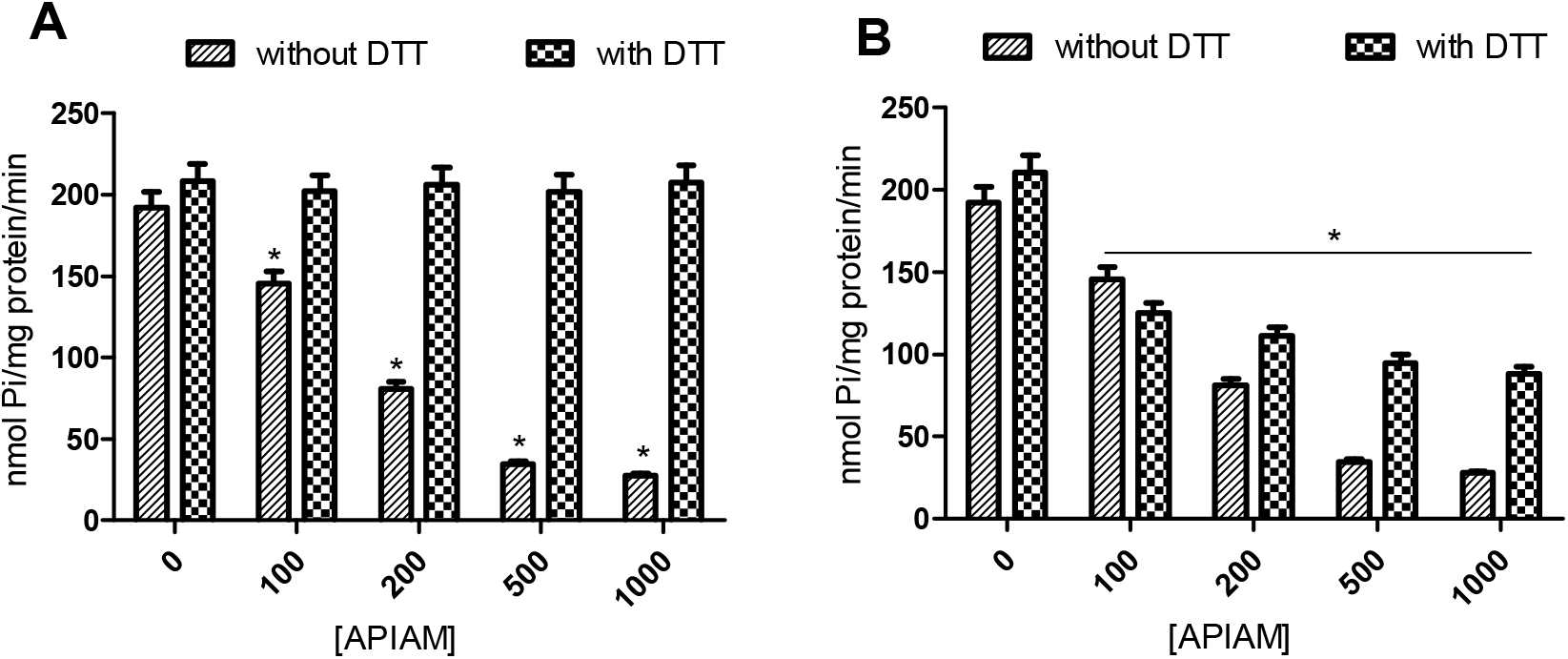
Effect of preincubation (panel A) and post-incubation (panel B) with 2mM DTT on APIAM-induced inhibition of the activity of the Na^+^/K^+^-ATPase at the ATP-binding site. Data were presented as mean ± SEM of at least three independent experiments carried out on different days. * indicates significantly lower than control (p < 0.05).

### Effect of IA and APIAM on Na^+^/K^+^-ATPase activity at the cation-binding sites

Previous submissions about the presence of thiol groups at the cation-binding sites of this protein necessitates investigation of the effect of these thiol modifying agents on the activity of the sodium pump at the cation-binding sites. Figure 4 showed (one-way ANOVA analysis) that while IA had no significant effect on the activity of the pump at all three cationic sites (panels A, C and E respectively), however, APIAM inhibited the activity of the pump significantly (p < 0.05), in a concentration-dependent fashion, when Na^+^, K^+^ and Mg^2+^ were excluded from the preincubating medium (panels B, D and F respectively).

**Figure 4:**
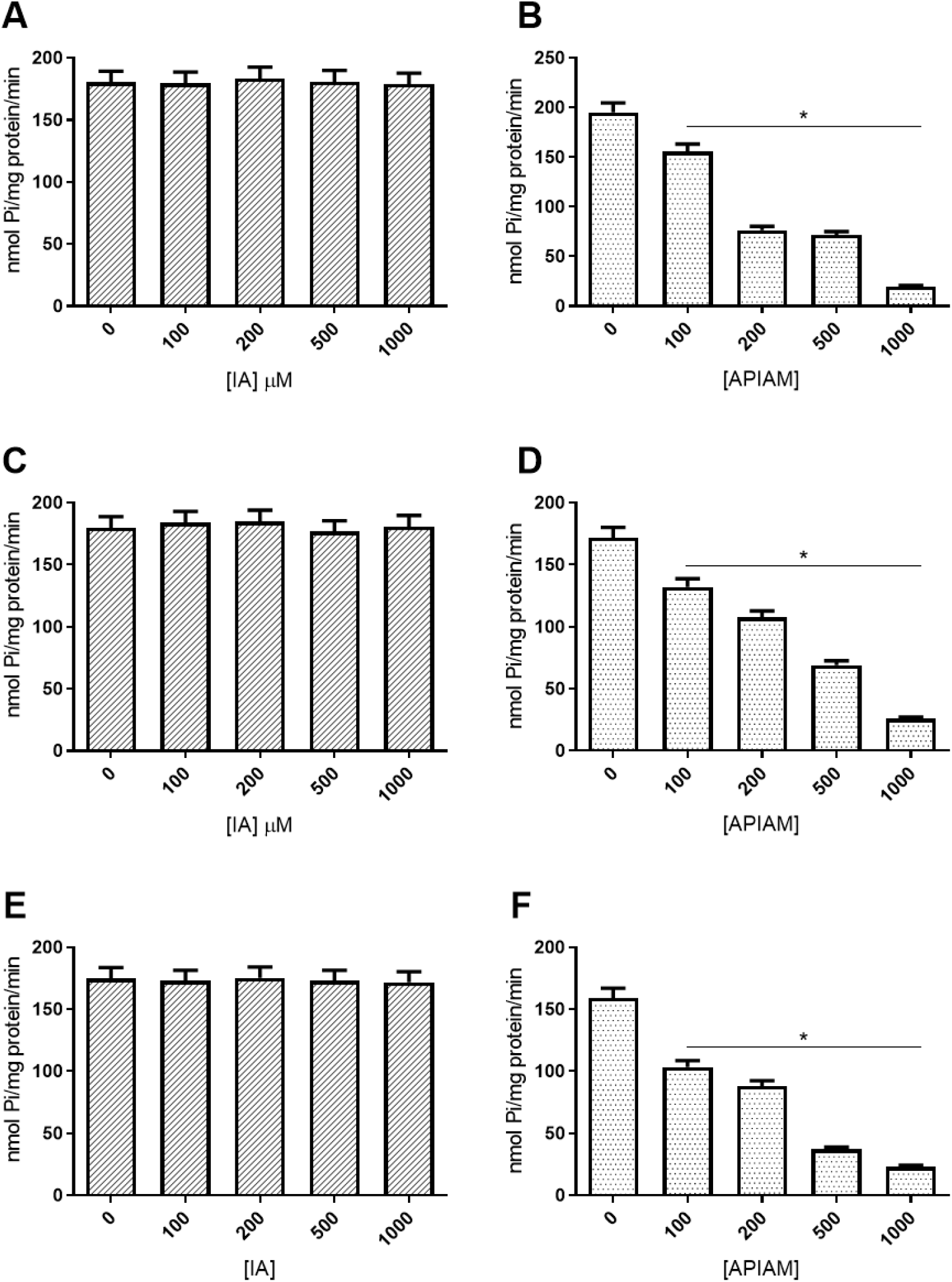
Effect of varying concentrations of IA and APIAM on the activity of the sodium pump at the Na^+^ (A and B respectively), K^+^ (C and D respectively) sites and upon exclusion of Mg^2+^ from the preincubation medium (E and F respectively). Data were presented as mean ± SEM of at least three independent experiments carried out on different days. # indicates significantly lower than control (p < 0.05).

### Effect of exogenous thiol, DTT, on APIAM-mediated enzyme inactivation at the cation-binding sites

Similar to the investigation of the nucleotide(ATP)-binding site, the effect of pre- and post-incubation with DTT on the enzyme inhibition mediated by APIAM on the activity of the pump when each of the cations were excluded from the preincubating medium was investigated. Two-way ANOVA analysis of the result as presented in Figure 5, shows that preincubation with DTT significantly prevented the inhibition induced by APIAM in the absence of each of the cations Na^+^ (panel A), K^+^ (panel C) and Mg^2+^ (panel E) from the preincubating medium (p < 0.05). However, post-incubation with DTT did not reverse the APIAM-mediated enzyme inactivation for all three cation-binding sites (panels B, D and F respectively).

**Figure 5:**
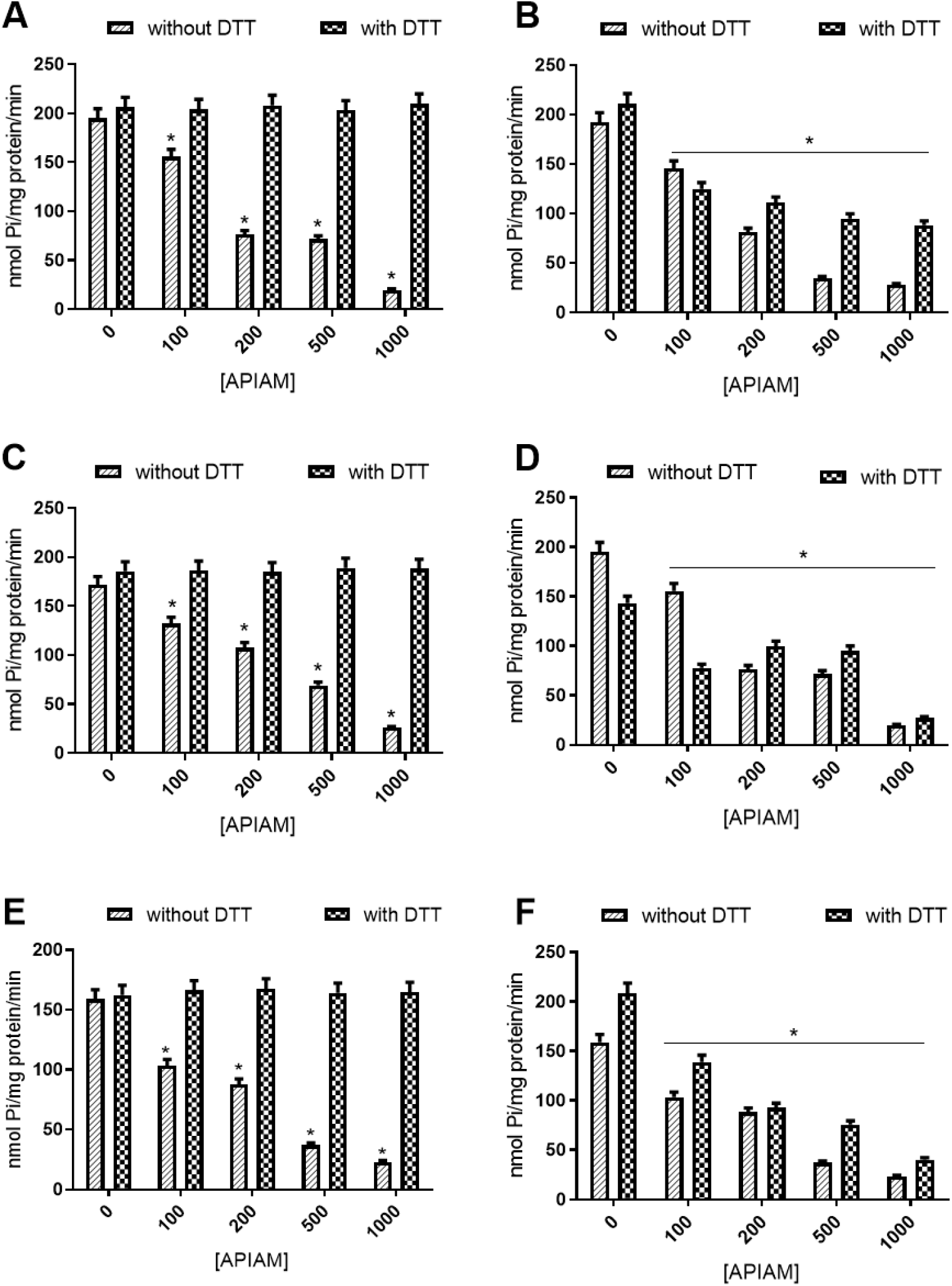
Effect of preincubation with 2mM DTT on APIAM-mediated inhibition of the Na^+^/K^+^-ATPase when Na^+^ (panel A), K^+^ (panel C), and Mg^2+^ (panel E) respectively were excluded from the preincubating medium, and the effect of post-incubation with DTT on APIAM-mediated inhibition upon exclusion of each of the respective cations from the preincubating medium (panels B, D and F respectively). Data were presented as mean ± SEM of at least three independent experiments carried out on different days. # indicates significantly lower than control (p < 0.05).

## DISCUSSION

The Na^+^/K^+^-ATPase is gaining attention as a target of oxidative assault as well as a toxin receptor in different neurodegenerative diseases (Renner et al., 2010; Petrushanko et al., 2016; DiChiara et al., 2017; Ohnishi et al., 2015; Shrivastava et al., 2015; Zhang et al., 2013). This transmembrane electrogenic enzyme is a redox-sensitive protein having well defined interactive domains specialized for its hydrolytic, catalytic, transport and signaling functions (Kaplan, 2002; Petrushanko et al., 2012, 2017, Kade et al., 2008). The redox sensitivity of this crucial enzyme is attributed to the presence of sulfhydryl groups at the interactive domains or sites which are highly vulnerable to oxidative modifications (Kade et al., 2008; Omotayo et al., 2011; 2015). Generally, the reactivity of protein thiols has been attributed to key factors such as (i) dissociation constant pKa of a thiol (ii) microenvironment (amino acid composition in vicinity to the cysteine residue), (iii) accessibility of the group and steric restrictions, and (iv) redox potential of a thiol (Nagy, 2013). Of primary importance to the present study is the microenvironment of these catalytically relevant thiols of the enzyme.

To appreciate the nature of the microenvironment of this enzyme, thiol modifying compounds, iodoacetamide (IA) and its derivative, N-acetyl-4-phenyliodoacetamide (APIAM) were employed. The first approach was to understand the reactivity of these compound with exogenous thiol compounds in the chemical model. Both compounds exhibited excellent thiol oxidizing property similar in mechanism since both possess the thiol reactive moiety present in the parent compound, iodoacetamide (IA). The mechanism of reaction highlights the ability of IA to modify thiol groups via bimolecular nucleophilic substitution reactions to form carbamidomethylated cysteines. This is a second order reaction that is dependent on the concentration of both the nucleophile (S_) and iodoacetamide (the substrate), as well as pH of the solvent (Wong and Liebler, 2008; Hill et al., 2009; Schmidt and Dringen, 2009).

The difference in the effect of these two thiol modifying agents on the activity of the Na^+^/K^+^-ATPase *in vitro* may not be unconnected to the presence of the acetyl-phenyl attachment in APIAM. The inability of iodoacetamide to inactivate the sulfhydryl enzyme (Figure 2, panel A), compared to its excellent thiol oxidizing potential in the chemical model, suggests a possible absence of interaction between iodoacetamide and the enzyme’s catalytically relevant thiols. The prevention of APIAM-mediated inhibition by preincubation with DTT (Figure 3, panel A) further confirms that the mechanism of enzyme inactivation is premised upon its thiol oxidizing property, a feature shared with IA in the chemical model. It is therefore apparent that the discrepancy is fundamentally based on the acetyl-phenyl attachment of APIAM. A similar phenomenon was also reported by Kade and coworkers, wherein two organoselenium compounds, dicholesteroyl diselenide (DCDSe) and diphenyl diselenide(DPDSe), exhibited a difference in their inhibitory effect on the activity of Na^+^/K^+^-ATPase. The authors suggested that the volume of the non-selenium group of DCDSe may pose an hinderance to its accessibility to the thiols at the catalytic site of the enzyme. This could be due to a number of factors, one of which is the microenvironment of the target protein thiols (Nagy, 2013). This feature has been defined by the pKa of distinct thiols, the amino acid composition in the vicinity of the cysteine residues as well as steric restrictions and localization of cysteine residue (Bogdanova et al., 2016; Nagy, 2013).

Moreover, the fact that the Na^+^/K^+^-ATPase is a transmembrane protein is very critical in the selective accessibility of its catalytically relevant sulfhydryl groups. This transmembrane protein experiences two distinct membrane-defined environments, a hydrophilic (extracellular/cytosolic) environment and a hydrophobic environment (within the membrane lipid bilayer) (Yoneda et al., 2020). Hence, it is likely that the presence of the acetyl-phenyl attachment which makes APIAM more lipophilic than IA facilitates its accessibility to these relevant thiols. Thus, we speculate that the catalytically relevant thiols at the nucleotide-binding domain of this enzyme are in a hydrophobic microenvironment.

Furthermore, earlier reports have demonstrated the presence of thiol groups at the cation-binding sites of this enzyme which are susceptible to oxidative modification (Kade et al., 2008; Omotayo et al., 2011). The observed inhibition of the activity of the enzyme at the Na^+^ and K^+^-binding sites (Figure 4) by APIAM and not IA further corroborates the above speculations about the microenvironment of the catalytically relevant thiol compounds at the nucleotide binding site. Apparently, the spatial localization of the thiol groups at the Na^+^ and K^+^-binding sites is in the hydrophobic core of the enzyme, possibly in the transmembrane domain, and not quite exposed to the aqueous environment. This could be nature’s protective design for these vulnerable thiol groups. More so, basal glutathionylation (dependent on redox status of the cell during protein folding) was reported for a number of cysteine residues in the alpha subunit of this enzyme, and described as a natural measure of preventing formation of disulphide bridges by cysteine residues in close proximity at the same cavity. This is because the formation of disulphide bridges between these cysteine residues leads to rigidity, and enzyme inactivation (Mitkevich et al., 2016).

To clarify that IA does not lose its thiol oxidizing property in the simple biological model *in vitro* used, the oxidation of protein-bound and free thiols was evaluated in the presence and absence of IA (data not shown). It was found that cytosolic free thiols are not the primary target of thiol modifiers. Thus, IA oxidized protein-bound thiols, speculatively, thiols unrelated to the enzyme’s transport function. It is thus clear that the inability of IA to inhibit enzyme activity is not due to a loss of its thiol oxidizing property but rather a question of accessibility to the microdomain of the catalytically relevant thiol groups. More so, in standard proteomic approaches, the use of IA for alkylation of protein thiols, did not provide complete alkylation of all thiols even after extended reaction duration (Galvani et al., 2001). This may also contribute to the observed inability of IA to inhibit the sulfhydryl Na^+^ /K^+^-ATPase.

Moreover, the binding of Mg^2+^ to ATP and formation of the MgATP^2-^ is essential for shielding it from the electronegative intracellular environment and supports the formation of the phosphate groups in the nucleotide (Jorgensen et al., 2003). Therefore, it is logical that the exclusion of Mg^2+^ from the reaction medium could induce a structural deformity in the ATP molecule, preventing its binding at the nucleotide-binding site and hence favour APIAM-mediated inhibition of the enzyme (Figure 4, panel F).

It is noteworthy that for the nucleotide and cation binding sites, while preincubation with DTT prevented enzyme inactivation by APIAM, post incubation with DTT is apparently ineffective in reversing enzyme inactivation (Figures 3 and 5). This phenomenon may be due to an irreversible modification mediated by APIAM binding. Previously, Petrushanko and collaborators identified basal glutathionylation of the Na^+^/K^+^-ATPase that is not reversible by DTT (Petrushanko et al., 2012). This observation was further attributed to posttranslational protein modification events related to the redox status of the cell (Mitkevich et al., 2016). While the present observation is not posttranslational, it can be inferred that the interaction of APIAM with thiols at these substrate binding sites could pose a steric hinderance to the intervention by DTT due to its acetyl-phenyl moiety. Further clarifications on these speculations are desirable.

Conclusively, the catalytically relevant thiols of the Na^+^/K^+^-ATPase are likely in an hydrophobic microenvironment which may be inaccessible to hydrophilic thiol oxidants. This appears to be a natural conservative and protective measure for the enzyme considering its essential physiological functions and necessary interaction with the aqueous intracellular and extracellular milieu. This exposes this enzyme to interactions with a diverse range of toxic agents. More efforts are being targeted at unveiling the details of the interactions of this enzyme with toxic oligomers as well as its specific inhibitors in the management cancer and neurodegenerative diseases. Further investigations on these findings hold much promise in unraveling the initiation and propagation of these diseases, paving way for their effective management.

## Acknowledgement

TIO and IJK acknowledge the financial support of DBT-India and TWAS. The authors also acknowledged the financial support of CAPES, FINEP, FAPERGS, PRONEX and CNPq, FINEP research grant Rede Instituto Brasileiro de Neurociencia (IBN-Net) # 01.06.0842-00 and the INCT for excitotoxicity and Neuroprotection-CNPq and CNPq-ProAfrica grant awarded to IJK and JBTR.

